# BrowseVCF: a web-based application and workflow to quickly prioritise disease-causative variants in VCF files

**DOI:** 10.1101/034769

**Authors:** Silvia Salatino, Varun Ramraj

**Author notes:** Corresponding author’s information: Name: Silvia Salatino.

## Abstract

As sequencing costs associated with fast advancing Next Generation Sequencing (NGS) technologies continue to decrease, variant discovery is becoming a more affordable and popular analysis method among research laboratories. Following variant calling and annotation, accurate variant filtering is a crucial step to extract meaningful biological information from sequencing data and to investigate disease etiology. However, the standard variant call file format (VCF) used to store this valuable information is not easy to handle without bioinformatics skills, thus preventing many investigators from directly analysing their data. Here, we present BrowseVCF, an easy-to-use stand-alone software that enables researchers to browse, query and filter millions of variants in a few seconds. Key features include the possibility to store intermediate search results, to query user-defined gene lists, to group samples for family or tumour/normal studies, to download a report of the filters applied, and to export the filtered variants in spreadsheet format. Additionally, BrowseVCF is suitable for any DNA variant analysis (exome, whole-genome and targeted sequencing), can be used also for non-diploid genomes, and is able to discriminate between Single Nucleotide Polymorphisms (SNPs), Insertions/Deletions (InDels), and Multiple Nucleotide Polymorphisms (MNPs). Owing to its portable implementation, BrowseVCF can be used either on personal computers or as part of automated analysis pipelines. The software can be initialised with a few clicks on any operating system without any special administrative or installation permissions. It is actively developed and maintained, and freely available for download from https://github.com/BSGOxford/BrowseVCF/releases/latest.

## Introduction

Recent developments of NGS technologies have led to a dramatic reduction in sequencing costs that, in turn, made DNA sequencing analyses accessible to small and medium-sized laboratories. Consequently, increasing amounts of sequencing data need to be analysed quickly and require specific bioinformatics expertise. Whilst large institutes might be able to rely on support from dedicated bioinformatics core units, wet-lab researchers from smaller centres may struggle to find easy-to-use tools for variant analysis. The variant call format (VCF), originally developed for the 1000 Genomes Project, has become the standard for storing DNA variants together with rich annotations (Danecek et al. 2011), and has led to the development of a variety of command-line analysis tools (for instance, VCFtools (Danecek et al. 2011) or GEMINI (Paila et al. 2013)). Recent efforts have been made towards developing more user-friendly graphical software to increase accessibility to non-bioinformaticians; examples include the commercial suites Ingenuity (http://www.ingenuity.com/), Alamut (http://www.interactive-biosoftware.com/alamut-visual/), GoldenHelix SNP (http://goldenhelix.com/SNP_Variation/index.html) and VariantStudio (http://www.illumina.com/informatics/research/biological-data-interpretation/variantstudio.html), as well as the open-source packages SNVerGUI (Wang et al. 2012), database.bio (Ou et al. 2015), gNOME (Lee et al. 2014) and BiERapp (Aleman et al. 2014). However, a disadvantage common to all these tools is that they strongly depend on a well-defined set of annotations, which limits the user to a restricted number of pre-defined features. Instead, researchers might be interested in keeping their personal or in-house annotations for downstream analysis, rather than discarding them. To our knowledge, only one recently-published software (VCF-Miner) addresses this problem by skipping the annotation step and focusing on the filtering part (Hart et al. 2015). However, its installation process requires administrative privileges and knowledge of virtual machines or Docker containers, it does not natively handle Variant Effect Predictor (VEP, http://www.ensembl.org/info/docs/tools/vep/index.html) annotations, and it does not allow the user to query variants that are annotated with a given set of genes of interest.

BrowseVCF overcomes these and other limitations, providing a faster, portable, and simpler interactive analysis tool to non-bioinformatician researchers. It has been optimised to reduce VCF loading and indexing time as much as possible by allowing the user to select only the fields of interest. The filter-history export feature was carefully designed to produce an easy-to-read report for posterity and reproducibility. It runs natively on multiple operating systems without the need for administrative privileges or knowledge of sophisticated deployment models. Moreover, our software has a new feature that allows users to perform keyword-based searches to identify, for instance, specific consequence terms or associated disorder states.

By applying filters sequentially to the input set of variants, BrowseVCF empowers researchers to narrow down the amount of putative variants to a small number of candidate disease-causing alleles.

## Results

### Benchmarks

We tested the performance of BrowseVCF on two different file types: an exome trio (proband, mother, father) and the whole-genome v2.18 of sample NA12878 generated by the "Genome in a Bottle Consortium” (https://sites.stanford.edu/abms/giab). As shown in Table 1, the amount of time required to pre-process the VCF file is below 2 minutes for standard exome data and the first query takes usually less than 30 seconds. Since each subsequent query uses the filtered output of the previous one, execution time gets shorter. For whole-genome data, pre-processing and index creation time is substantially longer than for an exome VCF (as expected). Timings were collected using a single core and differed between the Windows and the GNU/Linux machines. The bundled WinPython with Windows is not as well optimised as a natively-installed Python, so Windows performance can be increased by using a different installed Python (this also does not require administrator privileges). Specifying a higher number of CPUs to use in step 2 speeds up the process of creating the Wormtables for the fields of interest.

**Table 1.** Performances of BrowseVCF on exome and whole-genome data. Pre-processing times vary between operating system due to implementation differences intrinsic to Python.

| VCF file | Size<br>(MB) | Variants | Samples | Step 1: Pre-processing |  | Step 2: Index creation |  | Step 3: Filtering time<sup>3</sup> |  |
| --- | --- | --- | --- | --- | --- | --- | --- | --- | --- |
|  |  |  |  | time<sup>1</sup> |  | time<sup>2</sup> |  |  |  |
|  |  |  |  | Windows | GNU/Linux | Windows | GNU/Linux | Windows | GNU/Linux |
| Exome_trio.vcf.gz | 22 | 100014 | 3 | 0min 22sec | 0min 18sec | 1min 2sec | 0min 35sec | 0min 28sec | 0min 14sec |
| GIAB_v2.18.vcf.gz | 313 | 2915731 | 1 | 6min 29sec | 5min 8sec | 37min 53sec | 9min 11sec | 13min 9sec | 3min 38sec |

<sup>1</sup>Step required to convert the input VCF file to a format accepted by Wormtable
<sup>2</sup>Wormtables generated for the following fields: CHROM+POS, ID, REF+ALT, QUAL, FILTER
<sup>3</sup>Query executed on the FILTER field, keeping only PASS variants
Note: The "Windows" machine had 8 GB RAM and bundled, non-optimised WinPython; The "GNU/Linux" machine had 8 GB RAM and optimised system Python. Other modules and libraries are at identical versions between the two systems. All operations were performed using only 1 CPU.

## Discussion

The crucial step in variant analysis studies is an accurate prioritisation of the identified mutations, in order to pinpoint a small number of putative candidates that might be causal for a given disease or phenotype. BrowseVCF is a fast and intuitive interactive software that enables clinicians and researchers to browse and query their data without the need of any bioinformatics expertise. The major advantages of BrowseVCF with respect to similar tools are its clean interface and fast implementation, both particularly useful when analysing millions of variants from whole-genome sequencing data. Additionally, BrowseVCF is very straightforward to set up on any machine as it does not require administrator privileges on a personal machine to install and run. However, BrowseVCF shows a propensity for shorter execution times on GNU/Linux machines over Windows machines of similar specification due to operating-system specific differences in the included WinPython.

A limitation of existing tools is the need to load the entire VCF file, even though no more than a dozen annotations may typically be used for the analysis. In contrast, BrowseVCF allows the user to choose which annotations to process before running the filters and, if needed, to go back and load additional fields for further filtering steps. Considering that the number of samples and annotations in VCF files are the main drivers of the load time, being able to restrict the analysis only to the subset of truly relevant fields saves both computational time and disk space.

In addition, our software can be used both interactively and as part of computational pipelines. Furthermore, the ability to perform all the analysis locally is a considerable advantage when dealing with clinically-sensitive files. Although web-based interfaces are easier to operate without the need for command-line scripts, they can pose a security risk when dealing with sensitive patient data (Smedley and Robinson 2015). We designed BrowseVCF with flexibility in mind: it can run on a personal laptop without administrative privileges, as part of a HPC environment, or (in cases where patient data are available to an organisation behind its firewall) deployed centrally as a web service. In cases where data sensitivity is less of an issue, BrowseVCF is deployable as a public online web service. In conclusion, BrowseVCF can significantly improve efficiency in navigating large data sets to find candidate disease variants, thereby streamlining the clinical research process.

BrowseVCF is under active development and maintenance. At the time of writing, release version 2.5 is the latest and includes all the features discussed herein. Development and testing continue for implementation of new features, such as saving and resuming sessions, and global configuration options. However, the current version shows stability and robustness on multiple platforms and types of VCF annotations and is ready for deployment in a production environment.

## Methods

### VCF file loading and pre-processing

The VCF file format is very well-defined (Danecek et al. 2011). It consists of a header section, containing an arbitrary number of meta-information lines that start with the symbol ’#’, and of a data section, containing one line per variant, split into eight mandatory columns: chromosome (CHROM), 1-based starting position of the variant (POS), unique identifier, if existing (ID), reference allele in the genome (REF), alternative allele(s) of the variant (ALT), Phred-scaled quality score (QUAL), flag for passed/failed control checks (FILTER) and variant-specific annotations (INFO), which can be an unrestricted number of either flags (present/absent) or key-value pairs. VCF files containing one or more samples also include a ninth column (FORMAT), used to define the information enclosed in each subsequent column, and a genotype column for each sample, regarding the allele combination, the genotype read depth and other metrics.

The loading step within BrowseVCF is straightforward. The user selects a VCF file from his personal computer through a pop-up dialog box, which is then pre-processed. This step is able to correctly parse a variety of annotations, including those generated by VEP. BrowseVCF can accept both uncompressed and compressed (*.gz) VCF file types. The software identifies every annotation field present in the input VCF file and presents this list to the user, who can then select the fields of interest that will be used to filter the variants (Fig. 1). Additional fields can be selected at any time, if needed.

**Figure 1.**
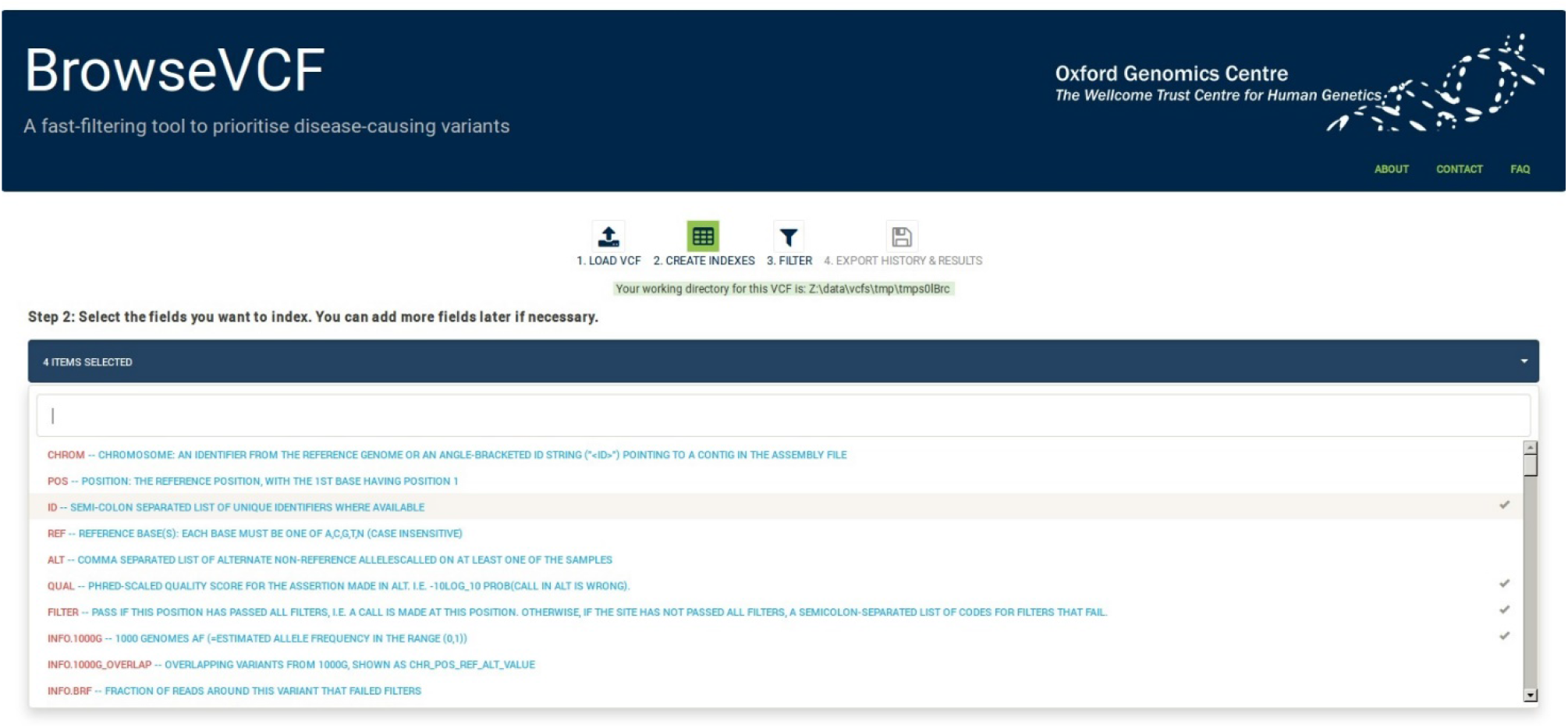
Screenshot of BrowseVCF step 2 (index creation). The drop-down menu lets the user browse through the annotation fields present in the input VCF file and select one or more fields to be used in filtering. A text box allows keyword searching across fields.

The “Wormtable” Python module (Kelleher et al. 2013) was chosen over other data storage systems for its efficient use of disk space and for the speed of its queries. However, as one Wormtable is created per chosen field, selecting fewer fields will reduce the time it takes for BrowseVCF to process the file. The pre-processing step is also required to convert missing values (’.’) to ’-1’ and to convert CSQ annotations (in case the file was annotated with VEP) into a Wormtable-compatible format.

### Variant prioritization by sequential filtering

Once the file is pre-processed, the user can start to filter variants according to five different query types (Fig. 2). The first filter lets the user query the data based on a given field of interest. Based on the type of annotation, different operators can be chosen for a specific cutoff/keyword (“greater_than”, “less_than”, “equal_to” or “contains_keyword”) and the user can choose whether to include or exclude variants with missing values. Moreover, the “contains_keyword” option allows searching for multiple keywords within the same field, which proves very useful when the objective is to keep variants labelled with different consequence or disease tags, for example. The second filter extracts variants with a given genotype in one, some or all samples; moreover, it is able to distinguish between “homozygous reference”, “homozygous alternate” and “heterozygous”. This feature is particularly useful in the study of inherited and *de novo* syndromes within a family or close relatives. BrowseVCF is able to process both diploid and non-diploid genomes alike. The third filter can retrieve all variants located within a given region of interest; in this case, the user is prompted to specify chromosome, start position and end position. This option is particularly useful to integrate other sources of information, e.g. regions identified by linkage analysis or targeted for capture. The fourth filter is able to extract either SNPs, InDels, or MNPs. Finally, the fifth query allows the user to specify a list of genes and retrieves all variants that are either associated (positive query) or not (negative query) to any of those entries within a specific annotation field. Filters are applied sequentially, so that a new filter is applied only to the variants that have successfully passed the previous filter.

**Figure 2.**
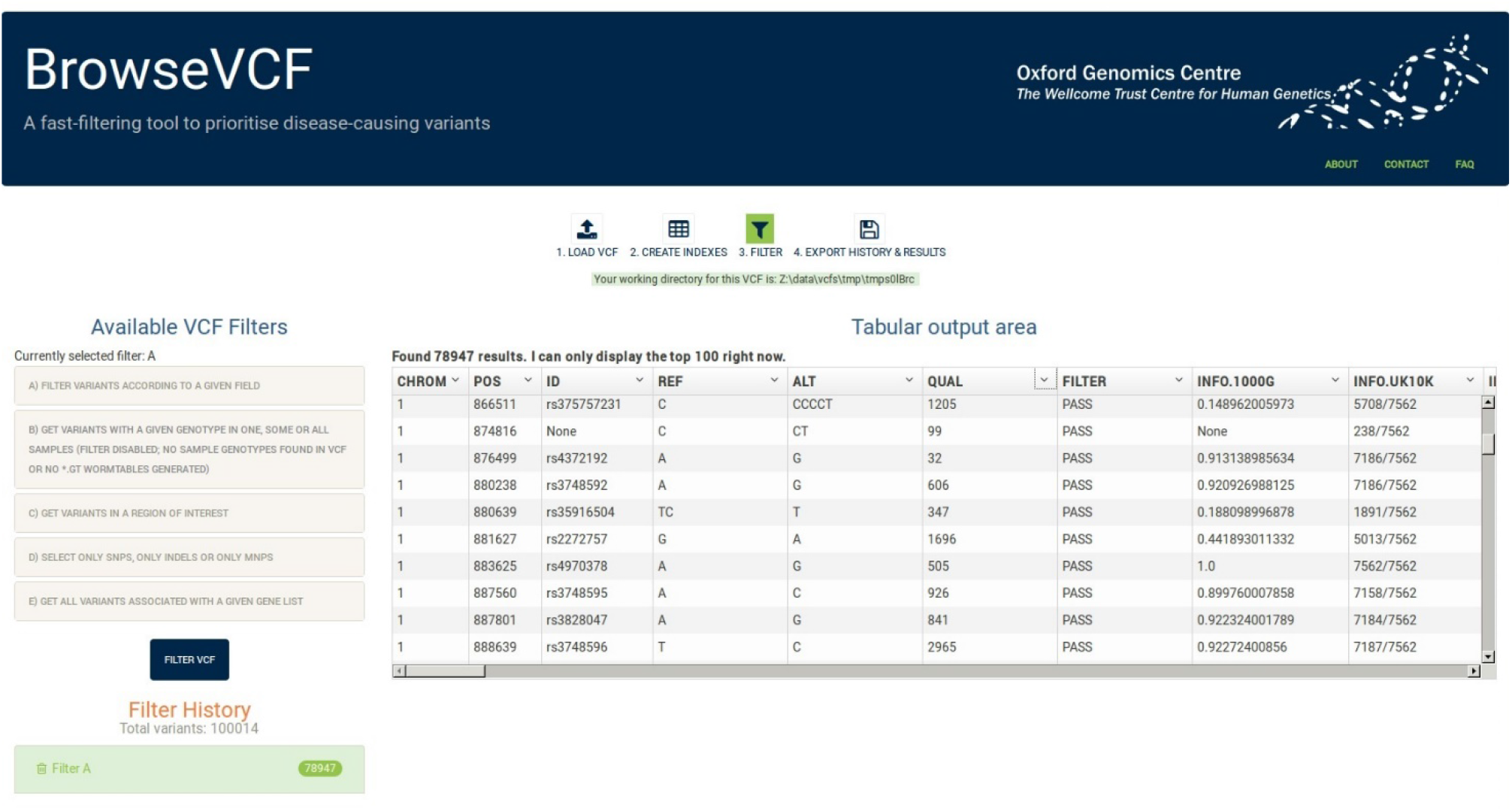
Screenshot of BrowseVCF step 3 (filtering). The left panel lists the five available filters, and expands to allow the user to define various filter options and cutoffs. The "Filter History” panel keeps track of all the sequential filters applied to the initial data and shows the output number of variants. The right panel is the output area, which displays the top 100 variants resulting from each consecutive filter. Fields (shown as columns) can be sorted or hidden if desired.

### Exporting results from the graphical interface

BrowseVCF comes with a user-friendly graphical interface (GUI) that allows researchers to load, filter, track, and export variants and annotations. Figure 2 displays the filtering interface. The right panel of the GUI displays variants in a tabular form, whereas the left panel keeps track of the filters applied and of the number of variants resulting from each query. The user can download the query history for record-keeping at any point. The output of each query is automatically stored to disk and can be deleted at any time from the "Filter History” section in the left panel. The right panel shows a preview of the top 100 variants and all variants can be exported at any time as a tab-separated text file, compatible with spreadsheet programs like Microsoft Excel.

### Technical details

BrowseVCF is free software under the GNU General Public License version 3 (http://www.gnu.org/licenses/gpl-3.0.en.html). It consists of a back-end service of Python scripts and a web server that runs locally, and of a front end built in JavaScript, CSS, and HTML5, leveraging popular cross-browser libraries such as AngularJS and Bootstrap. To perform fast and memory-efficient queries, all filtering operations are executed using Wormtable, a recently-developed Python module optimised to store genomic data in a compact binary tabular format through Berkeley DB, and retrieve data quickly by indexing desired fields (Kelleher et al. 2013). It is this use of Wormtable for data storage that makes BrowseVCF powerful, as future filters and enhancements can be implemented relatively easily by amending the indexes that are created in the Wormtable. The back-end scripts pre-process input VCFs and provide an interface to communicate with Wormtable in an intuitive manner, both on the command-line and through the web interface. To enable cross-platform usage, especially on Windows, Wormtable was altered and optimised. Every operation is carried out locally on the user’s computer, using the default browser; BrowseVCF is supported by all modern browsers, including Firefox and Chrome. The Windows version of BrowseVCF is shipped along with a lightweight distribution of WinPython, a bundled environment pre-configured to run directly on Windows. This enables the Windows user to simply double-click on an executable file without needing administrative permissions to install Python and necessary custom modules, which greatly improves the accessibility and ease-of-use of our software. However, since the tool can be configured to run with any web server back-end, it can also be installed site-wide within an organisation, thereby enabling shared use.

### Command-line version of BrowseVCF

BrowseVCF can also be used from command-line as a set of one-off queries, or as part of automated analysis pipelines. Each Python script is independent from the others and the list of required arguments can be displayed by typing the script name followed by "--help”. Some of the scripts are capable of parallel execution by specifying the number of cores at invocation, allowing BrowseVCF to operate within a high-performance computing (HPC) environment.

## Data access

BrowseVCF is freely available for download from https://github.com/BSGOxford/BrowseVCF/releases/latest.

## Acknowledgements

We thank Stefano Lise for advice and discussions, and Jerome Kelleher for significant help with implementation of new features in Wormtable. We also thank Kate Thomson, John Taylor, Hayley Mountford, and Alistair Pagnamenta for useful discussion, bug testing, and feedback on BrowseVCF, and Helen Lockstone for feedback on the manuscript. This work was funded by a Wellcome Trust Core Award (090532/Z/09/Z), supporting the Core facilities at The Wellcome Trust Centre for Human Genetics, including the Oxford Genomics Centre.

## Author contributions

S.S. designed the project. S.S. and V.R. implemented the code. S.S. and V.R. wrote the manuscript. All authors have read and approved the manuscript.

## References

Aleman A, Garcia-Garcia F, Salavert F, Medina I, Dopazo J. 2014. A web-based interactive framework to assist in the prioritization of disease candidate genes in whole-exome sequencing studies. Nucleic Acids Res 42: W88-93.

Danecek P, Auton A, Abecasis G, Albers CA, Banks E, DePristo MA, Handsaker RE, Lunter G, Marth GT, Sherry ST et al. 2011. The variant call format and VCFtools. Bioinformatics 27: 2156-2158.

Hart SN, Duffy P, Quest DJ, Hossain A, Meiners MA, Kocher JP. 2015. VCF-Miner: GUI-based application for mining variants and annotations stored in VCF files. Brief Bioinform doi:10.1093/bib/bbv051.

Kelleher J, Ness RW, Halligan DL. 2013. Processing genome scale tabular data with Wormtable. BMC Bioinformatics 14: 356.

Lee IH, Lee K, Hsing M, Choe Y, Park JH, Kim SH, Bohn JM, Neu MB, Hwang KB, Green RC et al. 2014. Prioritizing disease-linked variants, genes, and pathways with an interactive whole-genome analysis pipeline. Hum Mutat 35: 537-547.

Ou M, Ma R, Cheung J, Lo K, Yee P, Luo T, Chan TL, Au CH, Kwong A, Luo R et al. 2015. database.bio: a web application for interpreting human variations. Bioinformatics 31: 4035-4037.

Paila U, Chapman BA, Kirchner R, Quinlan AR. 2013. GEMINI: integrative exploration of genetic variation and genome annotations. PLoS Comput Biol 9: e1003153.

Smedley D, Robinson PN. 2015. Phenotype-driven strategies for exome prioritization of human Mendelian disease genes. Genome Med 7: 81.

Wang W, Hu W, Hou F, Hu P, Wei Z. 2012. SNVerGUI: a desktop tool for variant analysis of next-generation sequencing data. J Med Genet 49: 753-755.

